# Structure and conservation of amyloid spines from the *Candida albicans* Als5 adhesin including similarity to human LARKS

**DOI:** 10.1101/2021.11.29.470514

**Authors:** Nimrod Golan, Sergei Schwartz Perov, Meytal Landau, Peter N. Lipke

## Abstract

*Candida* Als family adhesins mediate adhesion to biological and abiotic substrates, as well as fungal cell aggregation and fungal-bacterial co-aggregation. The activity of at least two family members, Als5 and Als1, is dependent on amyloid-like protein aggregation that is initiated by shear force. Each Als adhesin has a ∼300-residue N-terminal Ig-like/invasin region. The following 108-residue, low complexity, threonine-rich (T) domain unfolds under shear to expose a critical amyloid-forming segment ^322^SNGIVIVATTRTV^334^ at the interface between the Ig-like/invasin domain 2 and the T domain of C*andida albicans* Als5. Amyloid prediction programs identified six potential amyloidogenic sequences in the Ig/invasin region and three others in the T domain of *C. albicans* Als5. Peptides derived from four of these sequences formed fibrils that bound thioflavin T, the amyloid indicator dye, and three of these revealed atomic-resolution structures of cross-β spines. These are the first atomic-level structures for fungal adhesins. One of these segments, from the T domain, revealed kinked β-sheets, similarly to LARKS (Low-complexity, Amyloid-like, Reversible, Kinked segments) found in human functional amyloids. Based on the cross-β structures in Als proteins, we use evolutionary arguments to identify functional amyloidogenic sequences in other fungal adhesins. Thus, cross-β structures are often involved in fungal pathogenesis and potentially in antifungal therapy.

**Importance:** Fungal adhesins form cell-to-cell bonds in biofilms. Many of the cellular interactions are dependent on formation of amyloid-like cross-β protein aggregates. Such structures are called ‘functional amyloids’ because they perform physiological activities and they are dependent on the same types of protein interactions that form the more familiar amyloid deposits in neurodegenerative diseases. We have identified sequence segments that form cross-β structures in the Als5 adhesin from the human pathogen *Candida albicans*. Such sequences are widespread among ALS family adhesins, including those from other human pathogens including *Candida auris*. Moreover, we revealed a structural similarity in a segment originating from Als5 threonine-rich low complexity region to human LARKS, pointing on a common structural motif coding for functional amyloids in different kingdoms of life.

## INTRODUCTION

Potential amyloid-like segments in fungal adhesins are functional in the sense that they mediate microbial adhesion, aggregation, and biofilm formation (1–4). They are amyloid in the sense that they mediate protein assembly into structured β-rich aggregates and fibrils, the same molecular interactions as those in amyloids in neurodegenerative diseases. Amyloid definition is based on the formation of cross-β fibrils composed of tightly mated β-sheets and interdigitating side chains (5–7). This structure leads to formation of birefringent structures that bind amyloidophilic dyes like Congo red and Thioflavins. Many microbial aggregates and biofilms display these dye-binding properties, and several adhesins have been demonstrated to form amyloid fibers *in vitro* (4, 8, 9). Among the best characterized, gram negative curli are pili composed of proteins assembled through cross-β amyloid-like interactions, and they mediate bacterial adhesion and biofilm formation. Like classical amyloids, biofilm formation can be inhibited by anti-amyloid compounds (9, 10). Thus, the characteristics of microbial functional amyloid mirror those of the better-known pathological amyloid in neurodegenerative disease and serum amyloidoses.

Classical amyloids form fibers that have a characteristic cross-β pattern under X-ray diffraction (7). This pattern shows strong orthogonal reflections of β-strands and β-sheets, with the β-strands orthogonal to the fiber axis and the sheets parallel to the axis. The β-sheets are stacked, and at least one interface between β-sheets is composed of tightly packed interdigitated sidechains, interacting through van der Waals forces and anhydrous H-bonds. This close interdigitation and dry interface give rise to the name “steric zipper,” and the crystalline cross-β structures of short segments are called “amyloid spines”(7).

A second type of homologous crystalline assembly has been recently described: Low-complexity, Amyloid-like, Reversible, Kinked segments (LARKS)(11). These structures also have a cross-β like arrangement, but the β-strands are kinked, the sheet interactions are dependent on smaller interfaces, and the structures are less stable and more evanescent. These LARKs spines are common in low-complexity unstructured regions of proteins (11). Low complexity domains are regions in protein sequences that differ from the composition and complexity of most globular proteins. Such LARKS sequences are involved in RNA-dependent phase separations that influence cellular stress response ((11, 12). LARKS have not been previously reported in adhesins. Among the best studied fungal adhesins are the paralogs Als1, Als3, and Als5 from *Candida albicans* (4, 12). Als1 and Als3 are important for fungal virulence in mice, and Als5 can attenuate immune response on the host (13, 14). Als adhesins are covalently attached to the cell wall glycans and are encoded at 8 loci in the genome (15, 16). They are typically 1200-2200 residues long (15). The N-terminal region is distal to the wall and consists of a 300-residue β-sheet-rich Ig-like/invasin region with two domains with Ig-like Greek key folds stabilized by disulfide bonds (17, 18). In each Als protein, the Ig-like/invasin region is followed by a 108-residue Threonine-rich low-complexity domain (T domain) that is highly conserved in all paralogs and includes a strong amyloidogenic sequence GIVIVA (19). C-terminal to T domains are 5-30 copies of a 36-residue tandem repeat. This region mediates hydrophobic effect interactions with abiotic substrates and with homologous structures. The C-terminal 500-1000 residues of Als proteins are also low complexity and are rich in O-glycosylated Ser and Thr residues, as well as in N-glycosylated Asn-Xaa-Ser/Thr motifs. This region forms a long flexible stalk that extends from the surface of the wall. The proteins are synthesized with C-terminal glycosyl phosphatidyl inositol (GPI) anchors, which are subsequently modified to form covalent crosslinks to cell wall glucans (20–22).

Als5 and other fungal adhesins have characteristics of functional amyloids. Under extension force applied by liquid flow or atomic force microscopy (AFM) extension, Als adhesin T domains unfold and expose the amyloidogenic sequence GIVIVA from the interface between the Ig/invasin region and the T domain (4). Peptides with this sequence form amyloid fibers, as do soluble forms of Als5, but peptides of fragments with a non-amyloid substitution GINIVA do not form amyloids (8, 19). When anchored to the cell surface, exposure of the GIVIVA sequence leads to formation of high avidity patches of adhesins on the cell surface. These patches consist of adhesins aggregated through amyloid-like bonds that are birefringent and bind amyloidophilic dyes. Recent evidence also shows that cell-to-cell binding is due to amyloid-like interactions between cells for Als5 and Als1 (23–25). Thus, formation of amyloid-like interactions mediates strong bonding between fungal cells, especially in biofilms formed under flow. Furthermore, amyloid-like structures coat the surface of fungi in fatal invasive fungal infections caused by many species. In superficial and invasive candidiasis, Als proteins are major components of these surface amyloid-like structures (14, 26). The fungal surface amyloid attenuates macrophage response to the invading fungi (13). Thus, extensive amyloid-like interactions are common on adhering fungi and in fungal biofilms and can influence host response.

Amyloidogenic sequences in fungal adhesins have been widely predicted, and force-dependent formation of amyloid-like surface nanodomains has been demonstrated in several cases (4, 27, 28). However, no atomic-level structures of amyloid spines or steric zippers from this type of protein have been reported. Therefore, we screened the Als5 Ig-like/invasin and T domains for predicted amyloidogenic sequences and determined structural characteristics of several of these. We have also analyzed the conservation and potential *in vivo* roles of these sequence segments in Als5 and homologous proteins.

## RESULTS

### Potential amyloid core sequences in *C. albicans* Als5

Sequence- and structure-based amyloid predictors use a variety of criteria including solubility and physicochemical properties, geometry, homology, and secondary structure propensity to predict occurrence of amyloidogenic core regions that can form cross-β aggregates and steric zippers. Accordingly, we scanned the primary structure of Als5 Ig/invasin and T domains (residues 20-433) with three amyloid prediction programs: TANGO, which has been useful for predictions in Als adhesins, is a thermodynamics state-based predictor (4, 29). AmylPred 2 is a consensus method based on eleven predictors (30). Lastly, FiSH Amyloid detects co-occurrence patterns in sequence data based on machine-learning techniques (31). Nine segments scored above threshold in at least two of the three predictors, marked on the Als5 sequence in Figure 1A. Six segments are in the Ig-like/invasin region: ^65^DTFILN^70^, ^132^IAFNVG^137^, ^156^NTVTFN^161^, ^168^SIAVNF^173^, ^196^IATLYV^201^, and ^259^FGISIT^264^, and three sequences are in the T domain: ^324^GIVIVA^329^, a section from the longer segment ^322^SNGIVIVATTRTV^334^ at the Ig/T domain interface, ^369^TSYVGV^374^, and ^388^TATVIV^393^.

**Figure 1.**
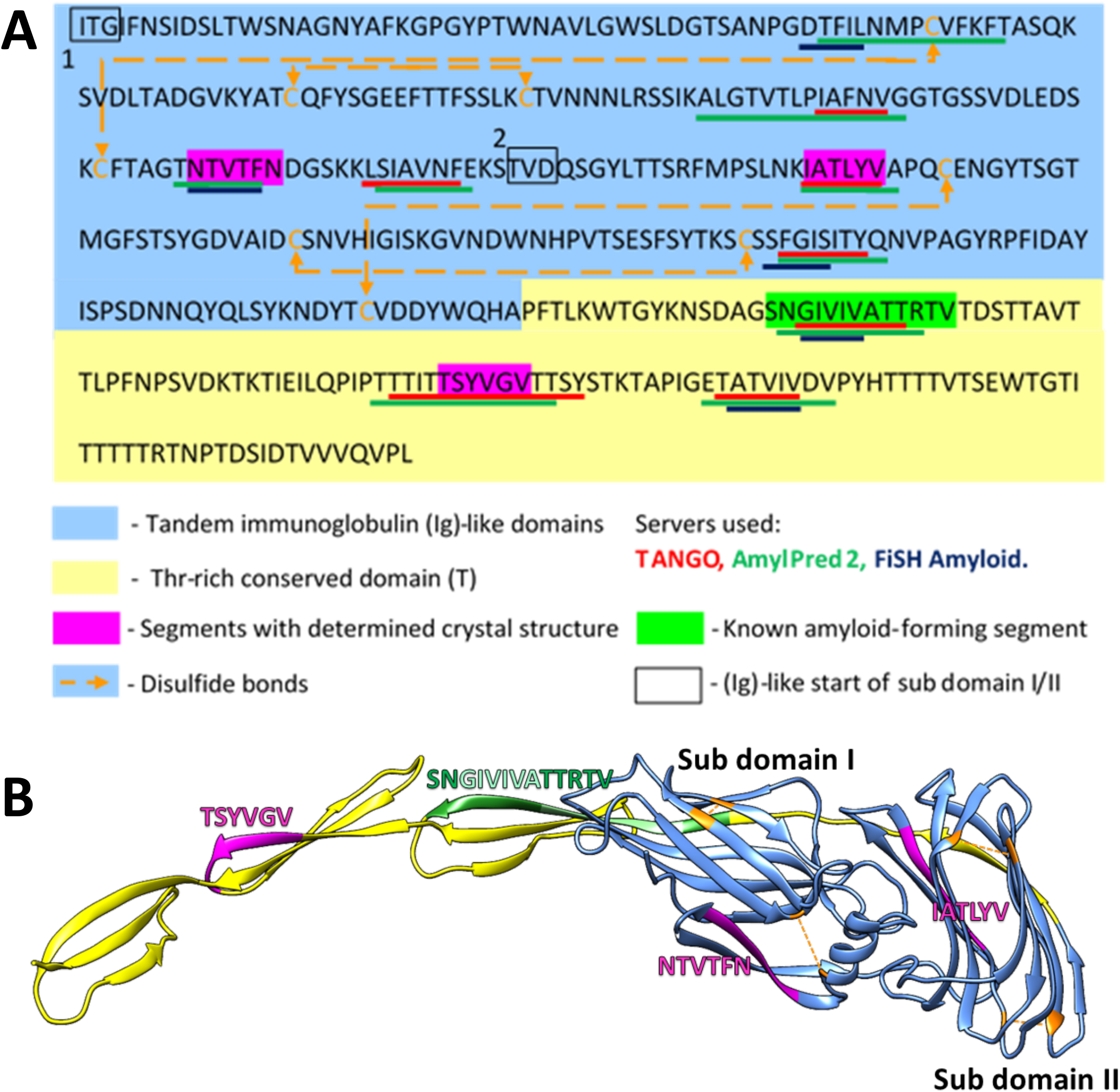
Predicted amyloidogenic segments within Als5^20-433^ sequence and modeled structure. (A) Annotated N-terminal residues 20-433 of the *Candida albicans* Als5 protein containing the ig-like/invasin and T domains, indicated by blue and yellow backgrounds, respectively. Segments with strong amyloidogenic predictions are denoted with colored bars under the sequence: TANGO (29) in red, AmylPred 2 (30) in green, and FiSH Amyloid in blue (31). Segments for which the crystal structure was determined are marked magenta, and the known amyloid segment ^303^SNGIVIVATTRTV^315^is marked green. Disulfide bonds are illustrated by yellow dashed arrow lines. (B) A three-dimensional (3D) model of the Als5 (residues 20-433) predicted by AlphaFold2 (32). The Ig/invasin region is based on homologous structures for Als3 with the two Ig subdomains indicated, the T domain is an *ab initio* model. Coloring schemes are as in panel A.

### Structures of amyloidogenic spines from Als5

We determined the atomic structures of ^156^NTVTFN^161^, ^196^IATLYV^201^ and ^369^TSYVGV^374^ segments (PDB IDs 6RHA, 6RHB, and 6RHD, respectively; Figure 2, Table 1 and Supplemental Table S1). The locations of the amyloid segments on the AlphaFold2 (32) predicted structure of Als5 are shown in Figure 1B. The crystal structure of the ^156^NTVTFN^161^ spine segment from the Ig-like domain formed a canonical steric-zipper structure of tightly mated parallel β-sheets with a dry interface. The ^196^IATLYV^201^ spine segment, also from the Ig-like domain 2, revealed a partial mating between antiparallel β-sheets. The ^369^TSYVGV^374^ spine segment from the T domain of Als5 exhibited an atypical amyloid structure, with antiparallel β-sheets and kinked β-strands (Fig. 2 and Supplemental Fig S1).

**Figure 2.**
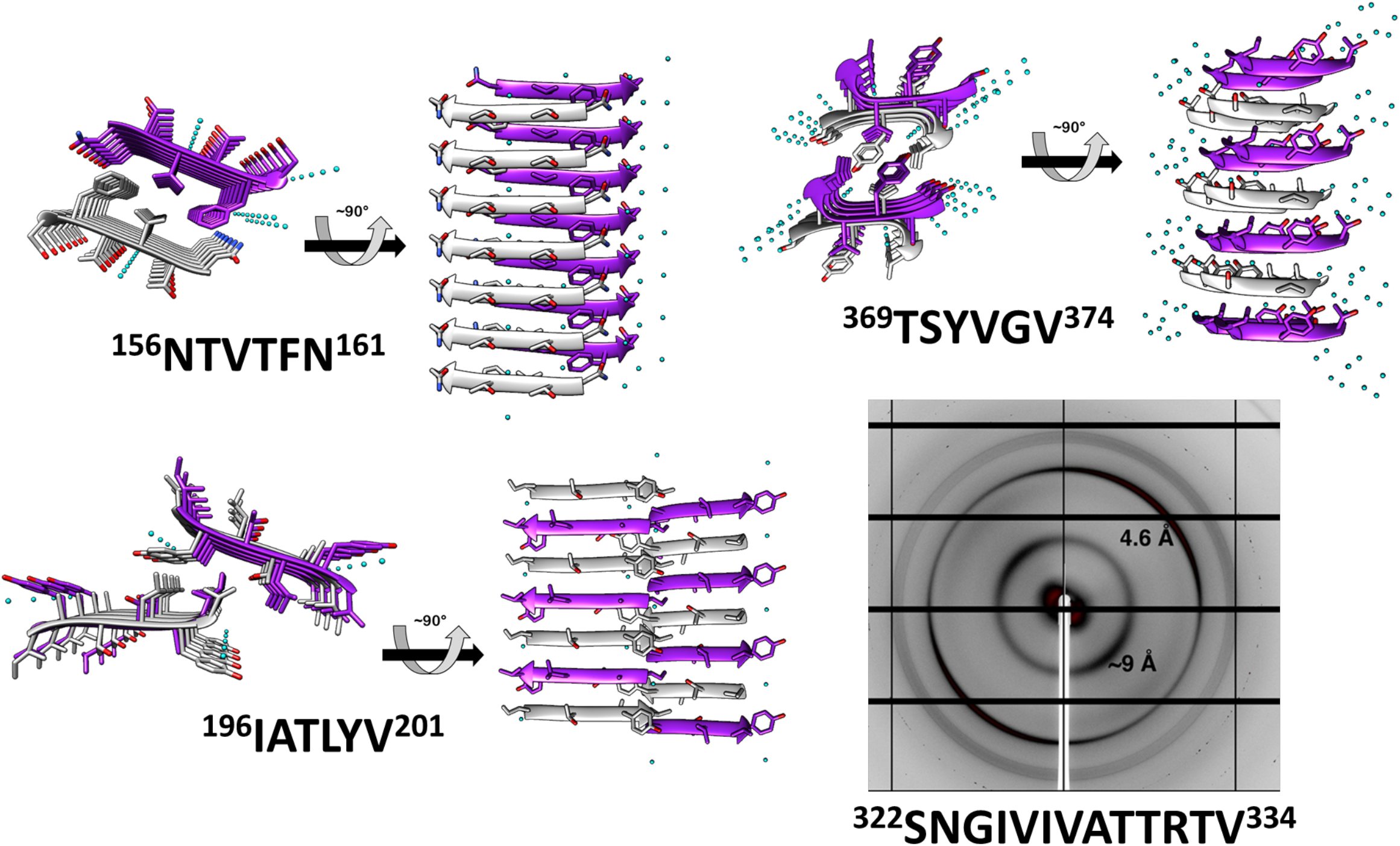
Crystal structures and fiber diffraction of Als5 amyloidogenic segments. High-resolution crystal structure of three predicted amyloidogenic segments of Als5 and the known amyloid segment ^303^SNGIVIVATTRTV^315^. The left panel of each structure is viewed down the fibril axis, with β-strands shown as ribbons and residues as sticks, while the right panel is viewed perpendicular to the fibril axis displaying β-strands running horizontally. Fibrils contain thousands of layers of β-strands, but only seven layers are shown here. Segment backbones are colored either gray or purple; heteroatoms are colored according to their atom type (nitrogen in blue, oxygen in red) and water molecules in cyan. The X-ray fiber diffraction pattern with major reflections labeled showing a cross-β pattern with orthogonal reflections at 4.6 Å and ∼9 Å spacings. Some reflections originating from ice also appear.

**Table 1.** Data collection and refinement statistics.

|  | Als5<sub>196</sub>IATLYV<sub>201</sub> | Als5<sub>156</sub>NTVTFN<sub>161</sub> | Als5<sub>369</sub>TSYVGV<sub>373</sub> |
| --- | --- | --- | --- |
|  | 6RHB | 6RHA | 6RHD |
| Beamline | ESRF ID23 2 | ESRF ID23 2 | ESRF ID23 2 |
| Date | September 7<sup>th</sup> 2015 | May 10<sup>th</sup> 2015 | December 11<sup>th</sup> 2015 |
| PDB ID |  |  |  |
| **Data collection** |  |  |  |
| Space group | P 2<sub>1</sub> 2<sub>1</sub> 2<sub>1</sub> | P 1 2<sub>1</sub> 1 | P 1 2<sub>1</sub> 1 |
| **Cell dimensions** |  |  |  |
| a, b, c (Å) | 9.49 17.87 22.42 | 4.87 39.81 20.26 | 11.77 9.36 16.79 |
| $\alpha$, $\beta$, $\gamma$ (°) | 90.0 90.0 90.0 | 90.0 90.0 90.0 | 90.0 95.1 90.0 |
| Wavelength (Å) | 0.8729 | 0.8729 | 0.8729 |
| Resolution (Å) | 22.4- 1.26 (1.32-1.26) | 39.8-1.60 (1.69-1.60) | 16.73-1.20 (1.25-1.20) |
| R-factor observed (%) | 24.6 (75.7) | 27.0 (100.5) | 20.0 (85.5) |
| <sup>a</sup>R<sub>meas</sub> (%) | 25.8 (79.5) | 28.0 (108.4) | 20.4 (90.0) |
| *I* / $\sigma I$ | 6.8 (2.9) | 7.3 (2.2) | 13.7 (2.5) |
| Test set size [%], selection | 10, random | 10, random | 10, random |
| Total reflections | 12416 (984) | 14151 (893) | 31912 (1290) |
| Unique reflections | 1076 (90) | 991 (126) | 1178 (139) |
| Completeness (%) | 91.1 (58.8) | 98.2 (90.0) | 96.2 (97.9) |
| Redundancy | 11.5 (10.9) | 14.3 (7.1) | 27.1 (9.3) |
| <sup>b</sup>CC1/2 (%) | 99.5 (97.1) | 97.5 (72.7) | 99.9 (84.2) |
| **Refinement** |  |  |  |
| Resolution (Å) | 13.97-1.26 (1.41-1.26) | 19.91-1.60 (1.64-1.60) | 16.73-1.20 (1.23-1.20) |
| Completeness (%) | 91.11 (76.6) | 97.9 (77.9) | 96.2 (98.8) |
| <sup>c</sup>No. reflections | 968 (218) | 891 (54) | 1059 (75) |
| <sup>d</sup>R<sub>work</sub> (%) | 9.97 (17.5) | 19.8 (29.0) | 10.6 (21.7) |
| R<sub>free</sub> (%) | 10.39 (24.1) | 21.3 (22.1) | 12.6 (22.3) |
| No. atoms | 49 | 103 | 51 |
| Protein | 48 | Chain A: 49<br>Chain B: 49 | 44 |
| Water | 1 | 5 | 7 |
| **B-factors** |  |  |  |
| Protein | 4.8 | Chain A: 11.1<br>Chain B: 12.4 | 7.25 |
| Water | 7.2 | 19.4 | 24.1 |
| **R.m.s. deviations** |  |  |  |
| Bond lengths (Å) | 0.007 | 0.014 | 0.011 |
| Bond angles (°) | 1.560 | 1.634 | 1.545 |
| Clash score (67) | 0.00 | 0.00 | 0.00 |
| Molprobability score (67) | 0.5 | 0.5 | 0.5 |
| Molprobability percentile (67) | 100th percentile | 100th percentile | 100th percentile |
| Number of xtals used for scaling | One crystal, one data set | Two crystals, one data set from each | One crystal, six data sets |

Values in parentheses are for highest-resolution shell.
<sup>(a)</sup> R-meas is a redundancy-independent R-factor defined in Reference (47).
<sup>(b)</sup> CC<sub>1/2</sub> is percentage of correlation between intensities from random half-datasets (48) .
<sup>(c)</sup> Number of reflections corresponds to the working set.
<sup>(d)</sup> Rwork corresponds to working set

These three peptides induced thioflavin-T (ThT) fluorescence with a typical amyloid curve of lag time followed by rapid aggregation (Fig. 3A) and formed amyloid-like fibrils visualized by transmission electron microscopy (TEM) (Fig. 3B). The peptides ^156^NTVTFN^161^ and ^196^IATLYV^201^ each formed abundant and long fibrils, while ^369^TSYVGV^374^ formed abundant short plump bundled fibrils. The longer peptide ^322^SNGIVIVATTRTV^334^ segment, known to be critical for biological activity of the adhesin and crucial for fibrillation of soluble forms of Als5 protein, formed long and relatively straight fibrils that also induced ThT fluorescence (4). This sequence contains a strongly predicted amyloid core sequence ^324^GIVIVA^329^. Although we did not successfully crystallize either the 6-residue or the 13-residue segments, the X-ray fiber diffraction of ^322^SNGIVIVATTRTV^334^ showed a canonical amyloid cross-β pattern with orthogonal reflection arches at 4.6 Å and ∼9 Å (Fig. 2).

**Figure 3.**
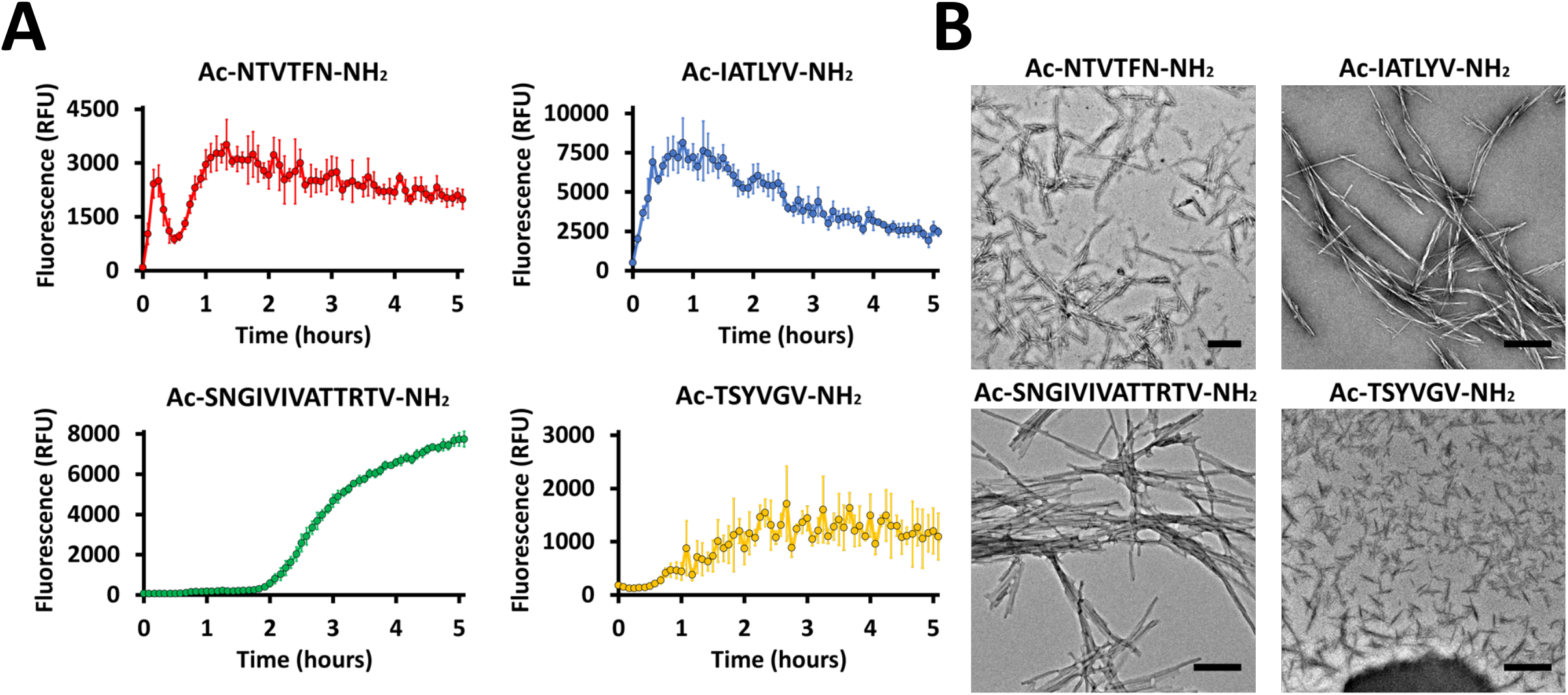
Predicted segments exhibiting amyloidogenic properties. (A) Segments for which the crystal structure was determined and the amyloid spines ^303^SNGIVIVATTRTV^315^ show fibrillation kinetics when incubated with ThT fluorescence dye. Measurements were conducted in 50 mM Tris-HCl buffer, 20 mM ThT, and 300 µM of peptide. The graph shows the average of three technical repeats, with error bars representing standard deviations. (B) Electron micrographs of the four segments show fibril formation after 48 hours of incubation. Scale bar represents 300 nm.

Because steric-zipper fibrils are unusual in that pairs of β-sheets mate more closely than the adjoining surfaces in other protein complexes, quantitative measures of amyloid stability are based on solvent-accessible surface area buried at the interface between the mating sheets, and shape complementarity indicating on the closeness of fit of two protein surfaces(33). The shape complementarity, inter-strand distance, and solvent-exposed surface area buried were calculated for Als5 spine structures and compared to the NNQQNY segment from yeast prion Sup35 steric zipper (PDB ID 1YJO) (34). The NNQQNY was chosen for this comparison as it shows one of the highest values of shape complementarity and surface area buried among steric zipper structures ((7)). The shape complementarity was similar for all Als5 spines segment, all lower than prion NNQQYY (Table S1). However, the T domain ^369^TSYVGV^374^ spine segment, displaying kinked β-strands, showed a significantly lower solvent-exposed surface area buried compared to the other structures (Table S1), indicating on a less stable structure that might also suggest reversible fibril formation.

The amyloid sequences are mapped onto an AlphaFold2 (32) based model of Als5 (Fig. 1B). The ^156^NTVTFN^161^ and ^196^IATLYV^201^ segments in the Ig-like/invasin region are each in close proximity to a disulfide bonded Cys residue (Figs. 1). Therefore, it is likely that these sequences would not have the conformational freedom to lead to solvent exposure or interaction necessary for amyloid formation. In contrast, the amyloid-forming segments in the T region, which is conformationally variable *in vivo*, are not constrained by disulfides or other structured regions. Thus, the amyloid propensity in this region is likely to be expressed *in vivo*, as demonstrated before for ^322^SNGIVIVATTRTV^334^ ((4).

### Patterns of evolutionary conservation and amyloidogenic propensities of amyloid segments in fungal adhesins

Evolution can shift and alter the functions of traits in all biological systems. Selection pressure will work to conserve traits that possess an important beneficial function. It is also true for functional amyloids, where purifying selection acts to conserve the amyloidogenic potential, amino acid sequence, and accessible conformation; however, the amyloidogenic potential and amino acid sequence are not always conserved equally. Occasionally, amino acid sequences can change considerably without greatly affecting their amyloidogenic potential (35). In light of this difference, we have examined the evolutionary conservation patterns of the amyloidogenic segments correlated with the trajectory of the change in amyloidogenic potential from close to more distant homologs of *C. albicans* Als5.

To evaluate the conservation of the amyloidogenic segments in protein homologous to Als5 we used the ConSurf webserver (36). First, we searched for homologs for Als5 residues 20-433 which include the Ig-like/invasin and T domains. Seventy-one unique sequences of homologs with 35–95% sequence identity were retrieved from the UniProt (UniRef90) database. We excluded homologs with large gaps in the multiple sequence alignment. The segments ^156^NTVTFN^161^, ^196^IATLYV^201^, ^322^SNGIVIVATTRTV^334^, ^324^GIVIVA^329^ and ^369^TSYVGV^374^ showed diverse pattern of sequence conservation across the Als5 homologs. In the Ig-like/invasin region, the sequence ^156^NTVTFN^161^ was remarkably conserved across the homologs (Fig. 4A), with five strongly conserved positions (Supplemental Fig S2), implying a conserved functional or structural role. In contrast, ^196^IATLYV^201^ was not well conserved and showed low average conservation value with four highly variable positions in its sequence. In the T domain, the ^322^SNGIVIVATTRTV^334^ region showed intermediate to above-average conservation. This observation is in agreement with previous reports showing that the functional amyloid-forming sequence ^322^SNGIVIVA^329^ is strongly conserved within the *C. albicans ALS* gene family (19, 25). The LARKS-like sequence ^369^TSYVGV^374^ also showed intermediate average conservation value with three highly conserved positions, with highest value at Gly^373^.

**Figure 4.**
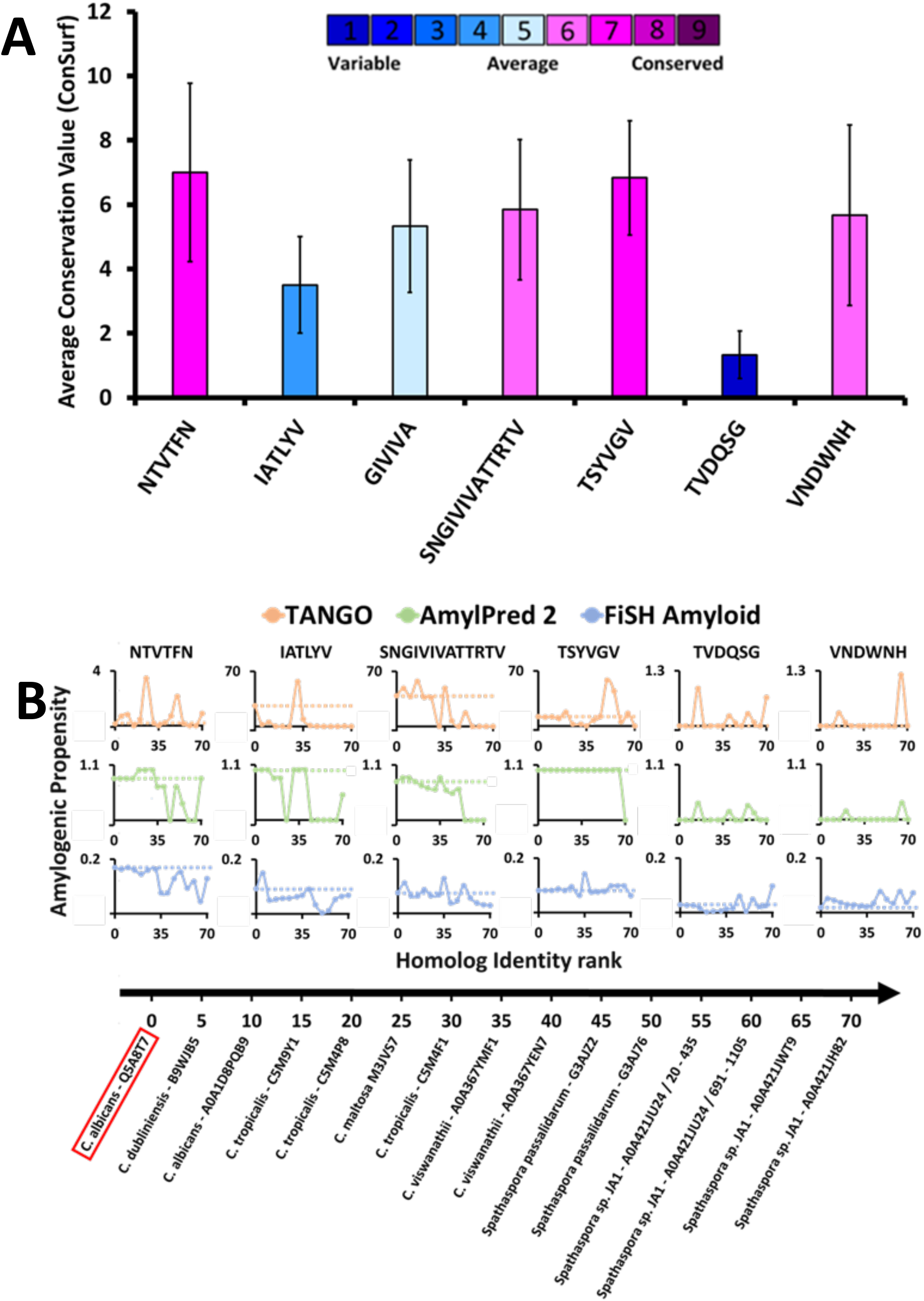
Amyloidogenic propensity and the evolutionary conservation of *C. albicans* Als5 segments. (A) Evolutionary conservation of Als5 segments. The graph displays evolutionary conservation values for seven segments (three structured segments, the amyloid spine ^303^SNGIVIVATTRTV^315^, and two control segments) from *C. albicans* Als5. These values were calculated based on a multiple sequence alignment of 71 homologs of Als5 generated using the ConSurf webserver ((36). The columns’ colors correspond to the average evolutionary conservation score (1 being the least conserved and 9 being the most conserved). Error bar represents the standard deviation of the average value of the residues in the sequence. (B) The amyloidogenic propensities of the above seven segments in Als5 orthologs (homologs from different species). Fourteen representative orthologs of Als5p were selected to represent a diverse group (as described in the Method section). The orthologs are plotted on the X-axis by their sequence similarity to *C. albicans* (evaluated based on ConSurf *e*-value parameter) from the most to the least similar homolog. The Y-axis shows the average amyloidogenic propensity of each Als5 segment and their equivalent regions in the fourteen orthologs. The amyloidogenic values were calculated using three servers and displayed using different colors: TANGO (orange) (29, 37) AmylPred2 (green) ((30), FiSH Amyloid (blue) (31). The different methods have a different score range, thus in each graph, a horizontal dashed line indicates the amyloidogenic value of the relevant segment from *C. albicans* Als5.

We also evaluated the conservation of amyloidogenic potential within the Als5p sequence homologs (Fig. 4B). The spine-forming segments ^156^NTVTFN^161^, ^196^IATLYV^201^, ^322^SNGIVIVATTRTV^334^, and ^369^TSYVGV^374^ presented a wide range of differences in amyloidogenic potential calculated by three predictors: TANGO, AmylPred 2, and FiSH Amyloid. For further analyses, we selected as controls two segments with a predicted low amyloidogenic propensity, ^177^TVDQSG^182^ and ^238^VNDWNH^243^, showing low and intermediate evolutionary conservation, respectively. The ^322^SNGIVIVATTRTV^334^ segment already implicated in amyloidogenic role of Als5 (4) showed a high amyloidogenic propensity in close homologs, which declined in more distant homologs. The LARKS-like sequence ^369^TSYVGV^374^, which is relatively conserved (Fig. 4A), shows the highest stability in maintaining amyloidogenic potential across homologs according to Amylpred2 and FiSH amyloid, with more fluctuations in propensity towards more distant homologs (Fig. 4B). In contrast, the Ig-like/invasin region segments ^156^NTVTFN^161^ and ^196^IATLYV^201^ show a less clear pattern of amyloidogenic propensity across homologs, similar to the control sequences ^177^TVDQSG^182^ and ^238^VNDWNH^243^. Control sequence ^177^TVDQSG^182^ showed high fluctuations in amyloidogenic propensity, despite its low value in the *C. albicans* proteins. This finding corresponds to its low sequence conservation (Fig. 4). Overall, the ^69^TSYVGV^374^ and ^322^SNGIVIVATTRTV^334^ segments in the Als5 T domain appear to have the most conserved amyloidogenic propensity among examined segments, at least across close homologs of *C. albicans*.

### Amyloidogenic propensity within the Als5 Ig-like/invasin and T domains in six species

We compared the amyloidogenic propensity of the entire Als5 Ig-like/invasin and T domains in Als5 homologs from seven species: *Candida dubliniensis, Candida tropicalis, Candida maltosa, Candida viswanathii, Spathaspora passalidarum, Spathaspora sp. JA1*, and *Candida auris* (39). The amyloidogenic propensities predicted by AmylPred2 are summarized in Figure 5. Each of the adhesins was predicted to have amyloid-forming sequence at the equivalent region to the ^322^SNGIVIVATTRTV^334^ segment of *C. albicans* Als5 (marked green in Fig. 5), except for one homolog from *Spathaspora sp. JA1*. The equivalent region to the LARKS-like segment ^369^TSYVGV^374^ also showed a strong amyloidogenic propensity (marked dark yellow in Fig. 5). In the *C. auris* homolog, there was no equivalent sequence to TSYVGV, but other segments in this region (residues 343-370) were predicted to be amyloidogenic (marked purple). In the Ig-like/invasin region, the equivalent sequence to ^196^IATLYV^201^ (marked blue in Fig. 5) showed no amyloidogenic propensity in *C. maltosa* and *S. passalidarum* or *Spathaspora sp. JA1*, but this segment did show some amyloidogenic propensity in *C. auris*. The predicted amyloidogenic propensity of this region was the least conserved among the distant homologs examined in Figure 4B. The second segment in the Ig-like/invasin region, equivalent to ^156^NTVTFN^161^, showed amyloidogenic propensity predictions (marked red in Fig. 5) across homologs except for *S. passalidarum* and *C. auris*. Overall, each ortholog showed in Figure 5 contains various regions with strong prediction for amyloidogenic propensity. Among the identified segments of Als5, we observed that the T-domain segments display more conserved amyloidogenic traits compared to segment in the Ig-like/invasin region.

**Figure 5.**
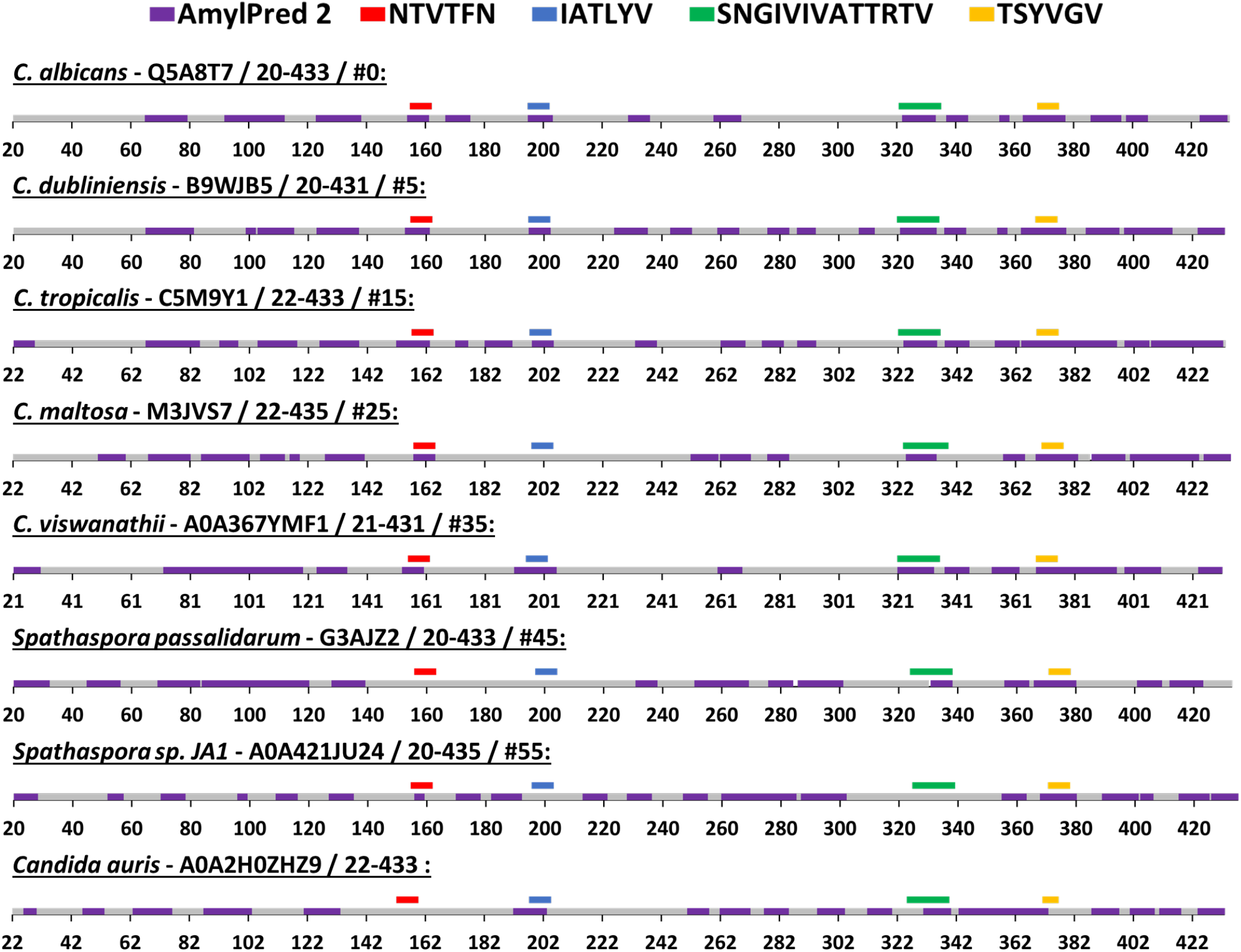
Predicted segments with amyloidogenic propensities in the Als5 Ig-like/invasin and T domains of *C. albicans* and seven orthologs. The amyloidogenic propensity, calculated using the AmylPred2 server (30), of Als5 Ig-like/invasin and T domains of *C. albicans* and seven orthologs. These specific orthologs were chosen as a diverse group among the 71 homologs collected by the ConSurf webserver along with *C. auris*, as described in the Method section. Each Ortholog is identified by the name of the species, their Swissprot ID, residue range of the Ig-like/invasin and T domains and their similarity to *C. albicans* ranked among 71 homologs (determined using the ConSurf-generated multiple sequence alignment). The regions with predicted amyloidogenic propensity are marked purple on the sequences, represented as a linear stretch with residue position marks. Regions equivalent to the *C. albicans* sequences for which the crystal structure was determined and the known amyloid segment ^303^SNGIVIVATTRTV^31^ are marked with different colors as indicated:^156^NTVTFN^161^ (red), ^196^IATLYV^201^ (blue), and ^369^TSYVGV^374^ (yellow) and ^303^SNGIVIVATTRTV^315^ (green).

## DISCUSSION

The structures and analyses presented here support the predictions of functional amyloid formation for fungal adhesins. Of the segments identified as potentially forming amyloids, we argue on the bases of structural and evolutionary constraints that two of them, in the T domain, are likely to function *in vivo*. We also showed atomic-level cross-β spine structures with known and novel geometries, including one LARKS-like structure reminiscent of functional and reversible human amyloids (11, 40).

### Amyloid predictions

Sequence- and structure-based amyloid predictors generally identify amyloidogenic or β-aggregation potential in many protein sequences. Indeed, systematic studies show that the majority of small peptides with high potential amyloidogenic sequences do form amyloids *in vitro* (7, 11, 41, 42). However, only few of the proteins containing these sequences form amyloids *in vivo*, probably because peptides are constrained by being part of a stable protein fold *in situ*. Formation of cross-β structures depends on conformation, solvent accessibility, and compatible geometry in flanking regions (5, 7, 42). Therefore, amyloids can often form only after proteins denature to expose the amyloidogenic sequences and allow flexibility in the peptide chains. In Als5, we identified 9 segments with high potential to form amyloid-like cross-β structures. We argue here two of these sequences are known to or may be involved in functional amyloid formation, while 6 others probably do not form cross-β structures *in situ* in the protein. The arguments are based on structural and evolutionary arguments, and on positive evidence for functionality of one segment.

### Potential amyloidogenic sequences in the Ig-like/invasin domain

Of the six potential amyloidogenic β-aggregating sequences in the Ig-like/invasin domain of Als5, three are within disulfide-stapled regions of the protein and so these segments are unlikely to be solvent-exposed and flexible enough to form cross-β structures (Figure 1). Of the three potential amyloidogenic sequences between that are not disulfide-constrained, ^156^NTVFN^160^ and ^196^IATLYV^201^ formed cross-β spines *in vitro*. However, these segments are unlikely to do so *in vivo.* Although the sequence ^156^NTVTFN^161^ is relatively highly evolutionary conserved (Fig. 4A and Supplemental Fig. S2), its TANGO amyloidogenic propensity is low and not conserved across homologs. It has greater potential in the other two predictors. (Figs. 4B and 5). In Ig/invasin domain II ^156^NTVTFN^161^ seals the edge of a β-sheet (Fig. 1), and so it contributes to the hydrophobic core of the fold. Therefore, this sequence is expected to be key for the high stability of this region, and this region of the protein unfolds only at high extension forces > 350 pN ((43, 44). In addition, ^156^NTVTFN^161^ is near disulfide-bonded Cys^150^ as well as the domain I/II interface, which both constrain its flexibility and geometry in the native protein. Therefore, this sequence is unlikely to form a cross-β structure *in vivo* in the intact protein.

The sequence ^196^IATLYV^201^ is in a β-strand (Fig. 1B) and is poorly conserved in sequence (Fig. 4A). Its equivalent region is predicted to have amyloidogenic propensity in some of the homologs, but without the specific trend of amyloidogenic propensity conservation along relatively close and distant homologs (Figs. 4B and 5). Therefore, it seems unlikely that this segment participates in the formation of functional β-aggregates in Als adhesins. The segment is extremely close to disulfide bonded Cys204, and so it is likely to be too geometrically constrained *in vivo* to form cross-β structures.

The four other potential amyloid sequences in the Ig-like/invasin region failed to form cross-β spines. Therefore, there is little support for formation of cross-β structures *in situ* within the Ig/invasin region of the protein.

### Potential amyloidogenic sequences in the T domain

Two of the potential amyloid sequences in the T domain show conservation of their sequence and amyloidogenic propensity. Figures 4B and 5 demonstrate the strong cross-β potential of ^322^SNGIVIVATTRTV^334^ and the conservation of this propensity in this region across homologs. Similarly, the LARKS-like segment ^369^TSYVGV^374^ also shows high conservation of amyloidogenic propensity (Figs. 4B and 5) and conservation in sequence as well (Fig. 4A and Supplemental Fig S2). Both of these segments showed strong cross-β properties (Fig. 3). Also contrasting with the Ig/Invasin domain, this region is unconstrained by disulfide bonds and becomes unstructured under low shear stress. Thus, these sequences are likely to be exposed and relatively unconstrained *in vivo* (4, 43, 45). These features are in accord with the known T domain functional amyloid formation.

### Functions of T domain amyloid sequences

The T domain and the sequence ^322^SNGIVIVATTRTV^334^ is essential for many activities of Als5 and its close paralog Als1 (4, 25). This sequence is necessary for strong cell-to-cell binding, as well as for clustering the adhesins on the cell surface. The cross-β nature of the interactions is supported by amyloid-like properties including birefringence, Congo red and thioflavin binding. A single site mutation V326N reduces the TANGO β-aggregation potential by about 20-fold, and reduces most aggregation activity in Als5 without affecting ligand binding to the Ig-like/invasin domain (9). Furthermore, adhesion activity is enhanced in the presence of a homologous peptide, and activity is reduced in the presence of a homologous non-amyloid peptide with the Val->Asn mutation. These are properties expected from a sequence interacting through β-aggregation. The T region unfolds easily under shear, including under physiological shear stresses (4, 28). Thus, extension forces unfold the T domain, rendering it unstructured giving the conformational flexibility needed for formation of cross-β structures.

The ^369^TSYVGV^374^ potential amyloid sequence is located ∼50 residues downstream in the T domain. This sequence should also be exposed when the domain unfolds under extension force (4, 43, 45, 46). It is conserved in sequence, and all homologs show similar amyloid-prone sequences at nearby positions (Figure 4A, Figure 5 yellow bars and Supplemental Figure S2). Under crystallization conditions, this peptide formed a structure like evanescent LARKS structures shown within low complexity regions (11). The conservation of amyloid potential implies that it may play a role in Als protein function, possibly to promote alignment of ^324^GIVIVA^329^ sequences in different proteins to facilitate formation of strong amyloid-like β-aggregates both *in cis* on a cell surface and *in trans* to form intercellular bonds. The third potential cross-β forming sequence in the T domain is ^388^TATVIV^393^, which is highly conserved in sequence and amyloid potential (Fig. 1 and Supplemental Fig. S2). However, we have no information on the relevance of this segment to Als5 function or to cross-β structures.

### Conclusions

Atomic resolution structures show that sequences in Als5 can form amyloid-like cross-β aggregates. Of the 4 sequences for which we have structural information, two in the Ig-like/invasin region have characteristics of fortuitous amyloid-formers, *i.e.* these segments have key roles in a hydrophobic domain core, coupled with propinquity to disulfide bonds and poor conservation of amyloid potential. In contrast, two sequences in the T domain show characteristics of force-dependent functional amyloids: conservation of sequence and amyloid potential, as well as demonstrated roles in activity of the protein for the ^324^GIVIVA^329^ sequence of Als5 and Als1 (4, 25). These results thus demonstrate a role for conservation of amyloid potential as well as sequence in determining functionality of naturally occurring sequences that form β-aggregates. Thus, evolutionary considerations for both sequence and amyloid potential are valuable tools to identify functional and pathological amyloid-forming sequences. Conversely, the same arguments can be used to identify many of the sequences that have high amyloid potential but nonetheless do not form amyloid structures *in vivo*.

## Materials and Methods

### Peptides and reagents

Peptide segments ^156^NTVTFN^161^, ^168^SIAVNF^173^, ^196^IATLYV^201^, ^324^GIVIVA^329^ and ^369^TSYVGV^374^ from Als5 (UniProt accession number Q5A8T7) were synthesized at >98% purity and purchased from GL Biochem (Shanghai) Ltd. The peptides were synthesized with unmodified termini for crystallography or with fully capped termini (acetylated in the N-terminus and amidated in the C structural terminus) for fibrillation assays. Thioflavin T (ThT) was purchased from Sigma-Aldrich. Dimethyl-sulfoxide (DMSO) was purchased from Merck. Ultra-pure water was purchased from Biological Industries.

### Computational prediction of amyloid spine segments

Amyloidogenic propensities of Als5 segments were predicted using combined information from several computational methods, including TANGO (29, 37), AmylPred 2 (30) and FiSH Amyloid (31). TANGO and FiSH Amyloid were used in their default settings, and AmylPred 2 was used with the default consensus option of only 4 of 9 predictors (TANGO and Amyloid Mutants were omitted from its prediction). We used TANGO, AmylPred 2, and FiSH Amyloid because they were the most sensitive and consistent predictors with quantifiable scores. The spine segments were defined as short segments of 6 residues, detected by at least two methods out of three at the epicenter of the predicted amyloidogenic area in the sequence (like in the case of ^369^TSYVGV^374^). Only in the case of the segment ^322^SNGIVIVATTRTV^334^, which is known to be critical to fibrillation of the Als5 protein (9), we worked with both its full length and a shorter version of 6 residues.

### Modelling of the Ig-like domain of Als5

The three-dimensional (3D) model was generated by AlphaFold2 (32) for the entire Als5 sequence from *Candida albicans*, taken directly from the UniProt accession number Q5A8T7. Using Chimera (51) we focused only on residues 20-433 which include the Ig-like/invasin and T domains. The per-residue confidence score (pLDDT) of the Ig-like domain ranged above 90 indicating a reliable prediction. For the T domain, the per-residue confidence score ranged between 70-90 indicating lower confidence in the prediction.

### Calculation of evolutionary conservation using the ConSurf/CenSeq webserver

We used ConSurf (36, 38) with default settings to examine the evolutionary conservation patterns of *Candida albicans* Als5 residues 20-433 which include the Ig-like and T domains. The calculation was run using ConSeq (52) (no structure was provided). One hundred and twenty-one unique sequences of homologs with 35-95% sequence identity were retrieved from the UniRef90 database. Homologs with large gaps (more than 5 residues) in their alignment were filtered using Jalview2 multiple sequence alignment editing program ((53). The alignment of the remaining 71 homologs was used for the calculations presented here. The alignment is accessible at <u>Link1</u>. The alignment of the seven orthologs used to generate the information presented in Fig. 5 is accessible at <u>Link2</u>.

### Amyloidogenic propensity correlated to the evolutionary conservation of the segments

Comparison between the segments ^156^NTVTFN^161^, ^196^IATLYV^201^, ^369^TSYVGV^374^, and ^322^SNGIVIVATTRTV^334^ evolutionary conservation was done by averaging the amino acid conservation scores attained from the ConSurf results for each segment. The segments ^177^TVDQSG^182^ and ^238^VNDWNH^243^ were taken as controls due to their lack of amyloidogenic propensity and their different conservation values, namely low and high respectively. The 71 retrieved homologs were ranked by their sequence similarity to *C. albicans* Als5, with the lowest *e*-value corresponding to the closer homolog to *C. albicans* Als5. Out of the 71 homologs we selected one along every five ranked homologs (in descending order of similarity to *C. albicans* Als5) for a total of fourteen sequences representing diverse sequences in the multiple sequence alignment. We then calculated the amyloidogenic propensity for each of the fourteen homologs using TANGO, AmylPred 2, and FiSH Amyloid (Figs 4B and 5). The multiple sequence alignment was used, via Jalview2 (53), to identify equivalent regions the segments of *C. albicans* discussed here. We then calculated the average predicted amyloidogenic propensity values in these corresponding sequences in each of the 14 homologs. TANGO averages were calculated by dividing the sum of each residue’s amyloidogenic propensity by its length. AmylPred 2 averages were calculated by assigning to each residue of the segments either a Boolean value of zero or one (based on consensus recognition of the method) and calculating the average similarly to TANGO averages. Using the default sliding window settings of the method, FiSH Amyloid averages were calculated by dividing the sum of the values assigned to all residues covering the segment by its length.

For the calculations of the amyloidogenic propensity of the entire Ig-like/invasin and T domains, we selected seven orthologs (homologs with a common ancestral parent in different species). Six of those were chosen along the ranked 71 homologs in gaps of 10 (starting from *C. dubliniensis* orthologs located five places away from the original *C. albicans* Als5). To these six, we added the emerging pathogen *Candida auris*, which was not a part of the homologs collected by the ConSurf webserver due to low sequence identity to the *C. albicans* Als5p (54, 55). The multiple sequence alignment of the 71 homologs along with the *C. auris* sequence was obtained by re-producing the ConSurf search with similar parameters but lower sequence identity of 25-95%, and then filtering out the irrelevant homologs using Jalview2.

#### Thioflavin T kinetic assays

Thioflavin T (ThT) is a commonly used for identifying and investigating the formation of amyloid fibrils *in vivo* and *in vitro*. In the presence of ThT, fibrillation curves commonly show delayed nucleation followed by rapid aggregation. Fibrillation kinetics of the peptides ^156^NTVTFN^161^, ^196^IATLYV^201^, ^369^TSYVGV^374^, and ^322^SNGIVIVATTRTV^334^ were monitored using ThT. All of the above peptides were synthesized with fully capped termini (acetylated in the N-terminus and amidated in the C structural terminus) to mimic its chemical nature in the full-length protein, and were dissolved to 10 mM in dimethyl sulfoxide (DMSO). Each of the freshly dissolved peptides was mixed with 50 mM Tris-HCl buffer with pH 7.3 and with 2 mM filtered ThT stock (made in ultra-pure water) to reach final concentrations of 300μM of peptide and 20 μM ThT in the final volume of 100 μl reaction. The reaction mixture was carried out in a black 96-well flat-bottom plate (Greiner bio-one) covered with a thermal seal film (EXCEL scientific) and incubated in a plate reader (CLARIOstar, BMG Labtech), at 37°C, with orbital shaking at 300 rpm, for 30 sec before each measurement. ThT fluorescence was recorded every two minutes using an excitation of 438±20 nm and an emission of 490±20 nm. The measurements were conducted in triplicates, and the entire experiment was repeated at least three times.

#### Transmission electron microscopy (TEM)

TEM was used to visualize fibrils. Samples for TEM were taken directly from the ThT kinetic assay plate, which was left to incubate at 37 C for 3 days with 300 rpm shaking in the plate reader. The TEM grids were prepared by applying 5 µl samples of each 300μM sample of peptide on 400 mesh copper grids with support films of Formvar/Carbon (Ted Pella), that were charged by high-voltage, alternating current glow-discharge (PELCO easiGlow, Ted Pella) immediately before use. Samples were allowed to adhere for 1 min followed by negative staining with 1% uranyl acetate for 1 min. Micrographs were recorded using a FEI Tecnai G2 T20 S-Twin transmission electron microscope at an accelerating voltage of 200 KeV at the MIKA electron microscopy center of the Department of Material Science & Engineering at the Technion.

### Fiber X-ray diffraction of the SNGIVIVATTRTV segment from Als5

The peptide SNGIVIVATTRTV was dissolved to 10 mg/ml in ultra-pure water. A few microliters were applied between two sealed glass capillaries until completely dried. X-ray diffraction of the samples was collected at the micro-focused beam P14 at the high brilliance 3rd Generation Synchrotron Radiation Source at DESY: PETRA III, Hamburg, Germany.

### Crystallization of Als5 segments

Peptides synthesized with free (unmodified) termini were used for crystallization experiments to facilitate crystal contacts. All peptides were dissolved to 10 mM in ultrapure water if possible, and in dimethyl sulfoxide (DMSO) in case they were water-insoluble. ^196^IATLYV^201^ was dissolved in 95% DMSO, ^156^NTVTFN^161 168^SIAVNF^173^ and ^324^GIVIVA^329^ were dissolved in 100% DMSO. ^369^TSYVGV^374^ was dissolved in 100% ultrapure water. Peptide solution drops (100 nL) were dispensed onto crystallization screening plates, using the Mosquito automated liquid dispensing robot (TTP Labtech, UK) located at the Technion Center for Structural Biology (TCSB). Crystallization using the hanging drop method, was performed in 96-well plates, with 100 µL solution in each well. The drop volumes were 150-300 nL. All plates were incubated in a Rock imager 1000 robot (Formulatrix), at 293 °K. Micro-crystals grew after few days and were mounted on glass needles glued to brass pins. No cryogenic protection was used. Crystals were kept at room temperature prior to data collection. Structures were obtained from drops that were a mixture of the following peptide and reservoir solutions: ^196^IATLYV^201^: 10 mM IATLYV, 0.1 M tri-Sodium citrate pH 5.6 and 1.0 M Ammonium phosphate. ^156^NTVTFN^161^: 10 mM NTVTFN, 0.1M HEPES 7.5pH, 0.8M NaH_2_PO_4_, 0.8M KH_2_PO_4. 369_TSYVGV^374^: 10 mM TSYVGV, 0.2M Sodium chloride, 0.1M Sodium acetate pH 4.6, 30% (v/v) 2-Methyl-2,4-pentanediol (MPD).

### Structure determination and refinement

X-ray diffraction data were collected at 100°K, using 5° oscillation. The X-ray diffraction data were collected at the micro-focus beamline ID23-EH2 of the European Synchrotron Radiation Facility (ESRF) in Grenoble, France; wavelength of data collection was 0.8729 Å. Data indexation, integration and scaling were performed using XDS/XSCALE (56). Molecular replacement solutions for all segments were obtained using the program Phaser within the CCP4 suite (57–59). The search models consisted of geometrically idealized β-strands. Crystallographic refinements were performed with the program Refmac5 (59). Model building was performed with Coot (60) and illustrated with Chimera (51). There were no residues that fell in the disallowed region of the Ramachandran plot. Crystallographic statistics are listed in Table 1.

### Calculations of structural properties

The Lawrence and Colman’s shape complementarity index (34) was used to calculate the shape complementarity between pairs of sheets forming the dry interface (Supplementary Table S1). The buried surface area was calculated with Chimera (UCSF), with a default probe radius and vertex density of 1.4 Å and 2.0/Å^2^, respectively. The number of solvent accessible buried surface areas was calculated as the average area buried of one strand within two β-sheets (total area buried from both sides is there double the reported number in (Supplementary Table S1).

## Acknowledgments

We acknowledge technical support provided by Yael Pazy-Benhar and Dikla Hiya at the Technion Center for Structural Biology (TCSB). We acknowledge support from Yaron Kauffmann from the MIKA electron microscopy center of the Department of Material Science & Engineering at the Technion, and Na’ama Koifman from the Russell Berrie Electron Microscopy Center of Soft Matter at the Technion, Israel. This research was supported by Israel Science Foundation (grant no. 2111/20), Israel Ministry of Science, Technology & Space (grant no. 78567), U.S.-Israel Binational Science Foundation (BSF) (grant no. 2017280), BioStruct-X, funded by FP7, and the iNEXT consortium of Instruct-ERIC. The synchrotron MX data collection experiments were performed at beamlines ID23-EH2 at the European Synchrotron Radiation Facility (ESRF), Grenoble, France, and at beamline P14, operated by EMBL Hamburg at the PETRA III storage ring (DESY, Hamburg, Germany). We are grateful to the teams at ESRF and EMBL Hamburg for their assistance.

## Author contributions

ML and PNL conceived the study. ML provided all support. All authors generated data, analyzed results, and contributed to writing and editing.

---

**Sup Figure 1.**
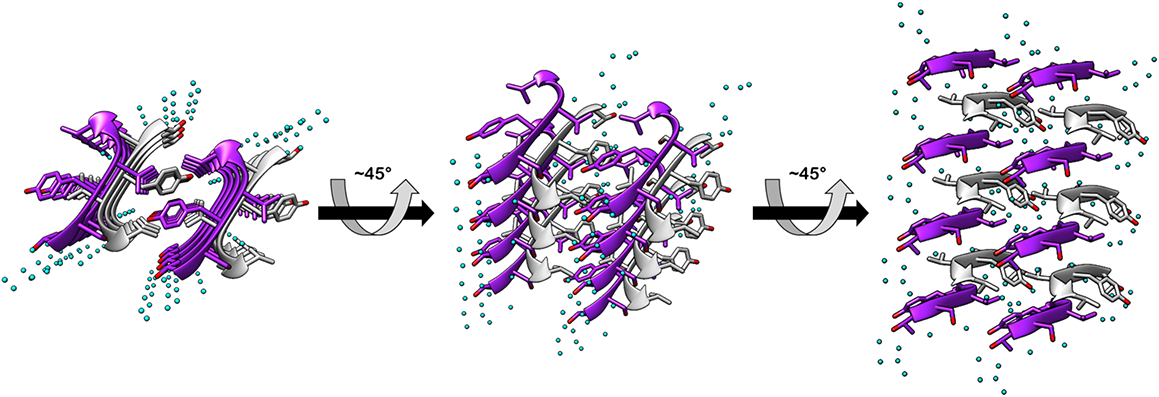
Crystal structures of the T-domain ^369^TSYVGV^374^ segments. The high-resolution crystal structure of ^369^TSYVGV^374^ is shown at different angles to better demonstrate its unique LARKS-like structure. The left panel is viewed down the fibril axis, with β-strands shown as ribbons and residues as sticks, while the middle and right panels respectively display a titled view (middle) and a view perpendicular to the fibril axis. The different view demonstrate the unique packed anti-parallel β-sheets composed of twisted strands, reminiscing the LARKS structures of kinked β-strands. β-sheet carbons are colored either gray or purple, and heteroatoms are colored according to their atom type (nitrogen in blue, oxygen in red) and water molecules in cyan.

**Supplemental Figure S2.**
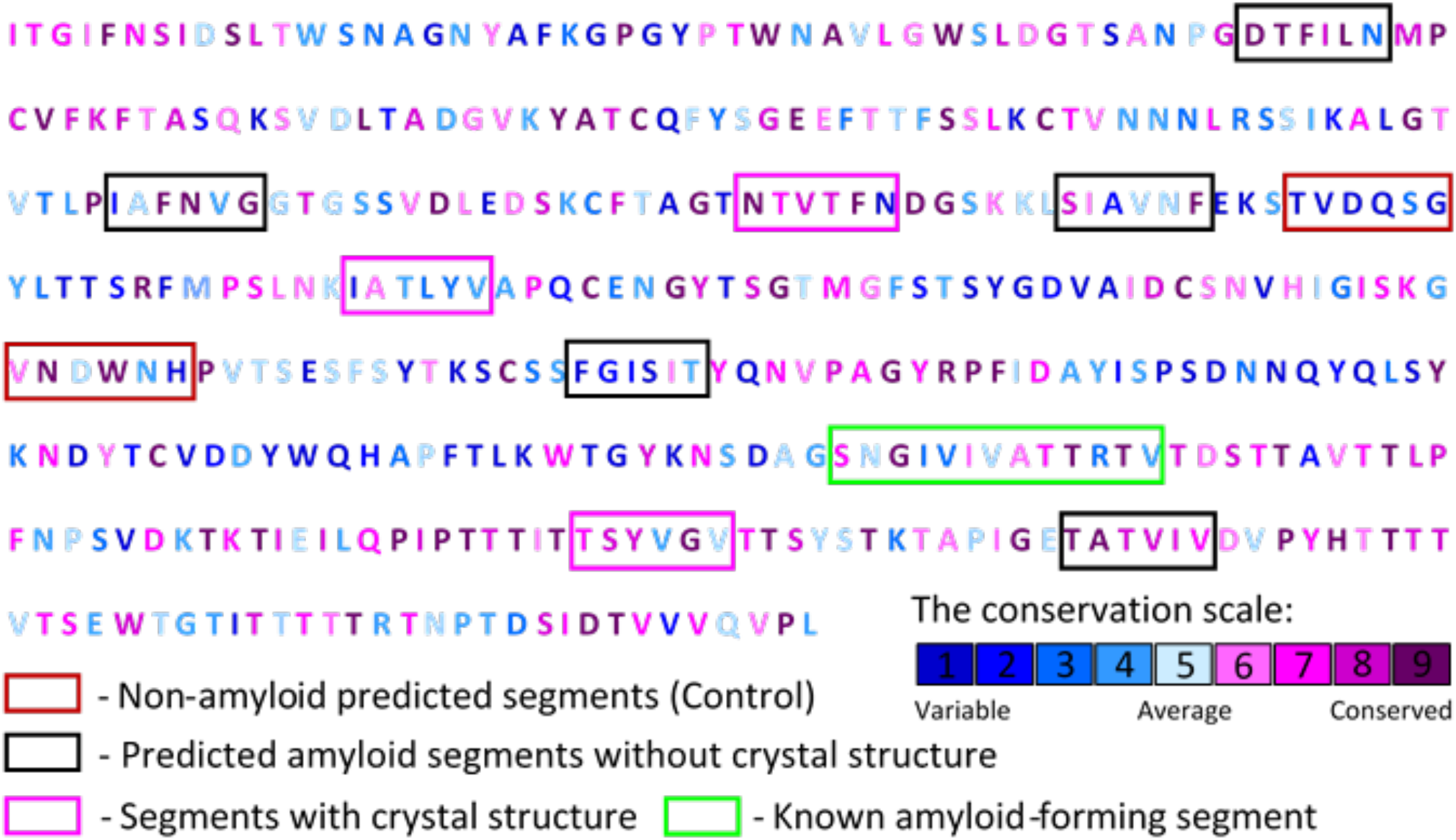
Sequence conservation scores mapped on the Als5(20-433) sequence. Evolutionary conservation scores were calculated using the ConSurf webserver (36, 38) based on a multiple sequence alignment and phylogenetic tree of 71 homologous proteins of the of N-terminal region of *Candida albicans* Als5 (UniProt ID Q5A8T7), residues 20-433. The conservation score of each residue is represented by a color scheme ranging from dark blue to dark purple (indicating variable to conserved). Segments for which the crystal structure was determined are marked with magenta boxes, the known amyloid segment ^303^SNGIVIVATTRTV^31^ is marked with a bright green box, and predicted amyloidogenic segments which didn’t crystalize are marked with black boxes. Segments that were not predicted to have amyloidogenic potential and used as control are marked with red boxes.

**Table S1.**
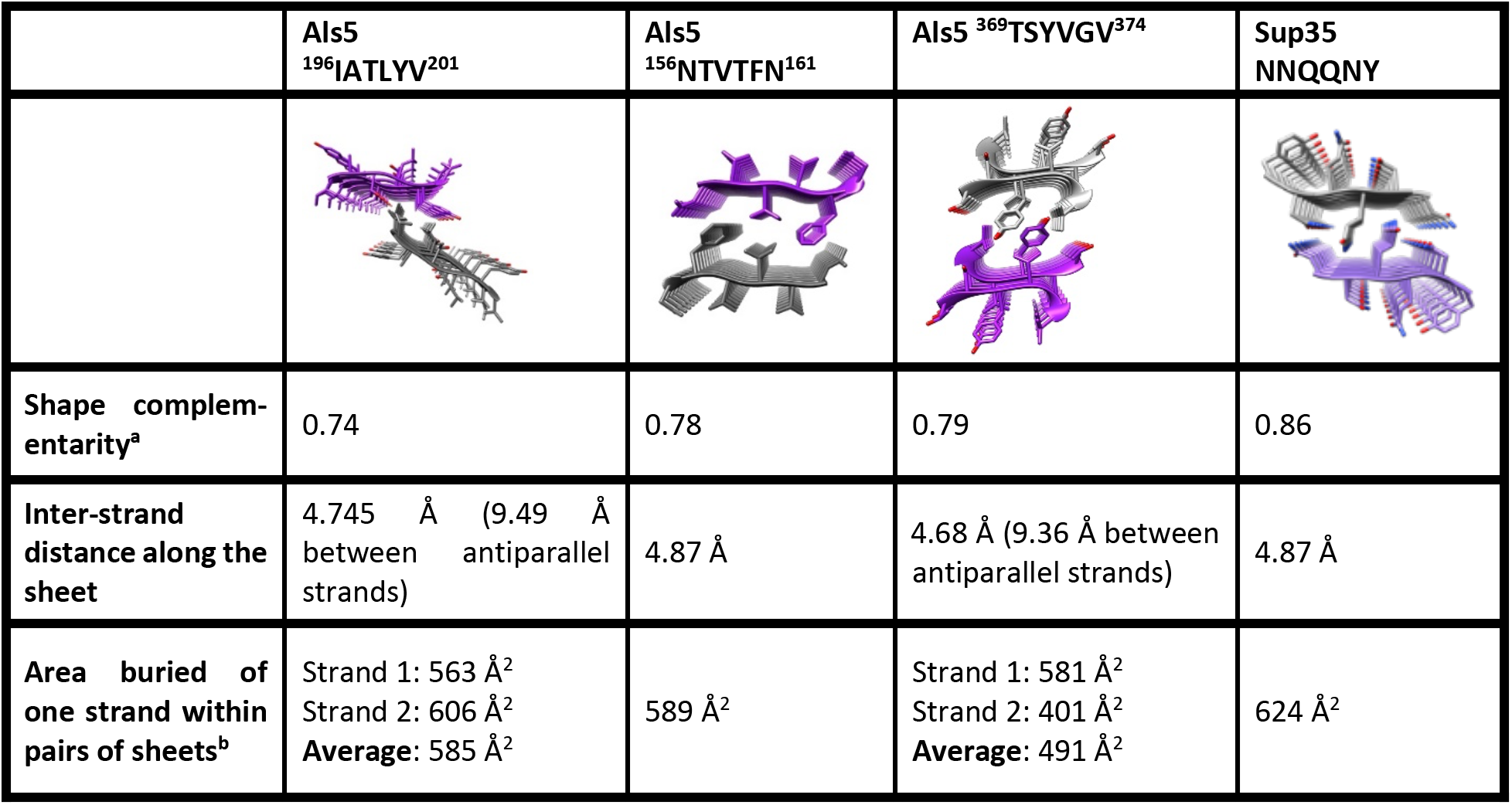
Features of the Als5 spine structures compared to the NNQQNY steric zipper structure. The values of shape complementarity, inter-strand distance and solvent exposed surface area buried calculated for the Als5 spine structures are compared with those of the NNQQNY segment from yeast prion Sup35 (PDB code 1YJO) (49) steric zipper structure. The NNQQNY was chosen for this comparison as it shows one of the highest values of shape complementarity and surface area buried among steric zipper structures (50). ^a^ Shape complementarity of 0 indicates no complementarity of the two surfaces and approaches 1 for atomic surfaces that fit perfectly together (34). ^b^ The solvent accessible surface area buried is the average area buried of one strand within two β-sheets (total area buried from both side is double the reported number). The surface area buried was calculated with Chimera (UCSF) (51) with default probe radius and vertex density are 1.4 Å and 2.0/Å^2^, respectively. IATLYV and TSYVGV assemble into antiparallel β-sheets, hence each of the antiparallel β-strands forms a difference interface that was calculated separately.

